# Reduction of Elevated Proton Leak Rejuvenates Mitochondria in the Aged Cardiomyocyte

**DOI:** 10.1101/2020.01.02.893362

**Authors:** Huiliang Zhang, Nathan N. Alder, Wang Wang, Hazel Szeto, David J. Marcinek, Peter S. Rabinovitch

## Abstract

Aging-associated diseases, including cardiac dysfunction, are increasingly common in the population. However, the mechanisms of physiologic aging in general, and cardiac aging in particular, remain poorly understood. Age-related heart impairment, especially diastolic dysfunction in HFpEF is lacking a clinically effective treatment. Using the model of naturally aging mice and rats, we show direct evidence of increased proton leak in the aged heart mitochondria. Moreover, we identified ANT1 as mediating the increased proton permeability of old cardiomyocytes. Most importantly, the tetra-peptide drug SS-31 prevents age-related excess proton entry, decreases the mitochondrial flash activity and mitochondrial permeability transition pore opening, rejuvenates mitochondrial function by direct association with ANT1 and the mitochondrial ATP synthasome, and leads to substantial reversal of diastolic dysfunction. Our results uncover the excessive proton leak as a novel mechanism of age-related cardiac dysfunction and elucidate how SS-31 is able to reverse this clinically important complication of cardiac aging.

## Introduction

Mitochondria are both the primary source of organismal energy and the major source of cellular reactive oxygen species (ROS) and oxidative stress during aging (Dai et al., 2014). Aged cardiac mitochondria are functionally changed in redox balance and are deficient in ATP production (Lesnefsky et al., 2016). Numerous reported studies have focused on redox stress and ROS production in aging (Dai et al., 2014). However, in its simplistic form, the free radical theory of aging has become severely challenged (Perez et al., 2009).

While more attention has been placed on mitochondrial electron leak and consequent free radical generation, proton leak is a highly significant aspect of mitochondrial energetics, as it accounts for more than 20% of oxygen consumption in the liver (Brand, 2005) and 35% to 50% of that in muscle in the resting state (Rolfe and Brand, 1996). There are two types of proton leak in the mitochondria: 1) constitutive, basal proton leak, and 2) inducible, regulated proton leak, including that mediated by uncoupling proteins (UCPs) (Divakaruni and Brand, 2011). In skeletal muscle, a majority of basal proton conductance has been attributed to adenine nucleotide translocase (ANT) (Brand et al., 2005). Although, aging-related increased mitochondrial proton leak was detected in the mouse heart, kidney and liver by indirect measurement of oxygen consumption in isolated mitochondria (Harper et al., 1998; Serviddio et al., 2007), direct evidence of functional impact remains to be further investigated. Moreover, the exact site and underlying mechanisms responsible for aging-related mitochondrial proton leak are unclear.

SS-31 (elamipretide), a tetrapeptide (D-Arg-2′,6′-dimethyltyrosine-Lys-Phe-NH2), binds to cardiolipin-containing membranes (Birk et al., 2013) and improves cristae curvature (Szeto, 2014). Prevention of cytochrome *c* peroxidase activity and release has been proposed as its major basis of activity (Szeto, 2014; Szeto and Birk, 2014). SS-31 is highly effective in increasing resistance to a broad range of diseases, including heart ischemia reperfusion injury (Cho et al., 2007; Szeto, 2008), heart failure (Dai et al., 2013), neurodegenerative disease (Yang et al., 2009) and metabolic syndrome (Anderson et al., 2009). In aged mice, SS-31 ameliorates kidney glomerulopathy (Sweetwyne et al., 2017) and brain oxidative stress (Hao et al., 2017) and has shown beneficial effects on skeletal muscle performance (Siegel et al., 2013). We have recently shown that administration of SS-31 to 24 month old mice for 8 weeks reverses the age-related decline in diastolic function, increasing the E/A from just above 1.0 to 1.22, restoring this parameter 35% towards that of young (5 month old) mice (Chiao et al., 2020). However how SS-31 benefits and protects aged cardiac cells remains unclear.

In this report we investigated the effect and underlying mechanism of action of SS-31 on aged cardiomyocytes, especially on the mitochondrial proton leak. Using the naturally aged rodent model we provided direct evidence of increased proton leak as the primary energetic change in aged mitochondria. We further show that the inner membrane protein ANT1 mediates the augmented proton entry in the old mitochondria. Most significantly, we demonstrate that SS-31 prevents the excessive proton entry and rejuvenates mitochondrial function through direct association with ANT1 and stabilization of the ATP synthasome.

## Results

### SS-31 alleviates the excessive mitochondrial proton leak in old cardiomyocytes

To examine whether SS-31 restores aging mitochondrial function, we applied the Seahorse mitochondrial stress assay to intact primary cardiomyocytes. The Seahorse Assay revealed higher mitochondrial basal respiration in cells from old mice than that in young mouse cells (Fig 1A, C); however, the maximal respiratory rate was not significantly different (Fig 1A, D). The increased basal respiration was attributable to a higher proton leak in old cardiomyocytes (164 ± 16 in 24 month vs 82 ± 12 in young, pmol/min/800cells, n=7-14, p<0.01) (Fig. 1A, B). Although, SS-31 has only a minor and non-significant effect on young cardiomyocytes (Fig. S1), acute *in vitro* treatment of isolated old cardiomyocytes with SS-31 (100 nM, 1 μM, or 10 μM for 2 hrs), caused reduced mitochondrial proton leak (Fig. 1A, B and Fig. S2), shifting their respiratory pattern to a more youthful state. These results indicate that SS-31 directly protects aging cardiac energetics through rapid rejuvenation of mitochondrial respiration in cardiomyocytes, and in particular, by reducing proton leak.

**Fig. 1:**
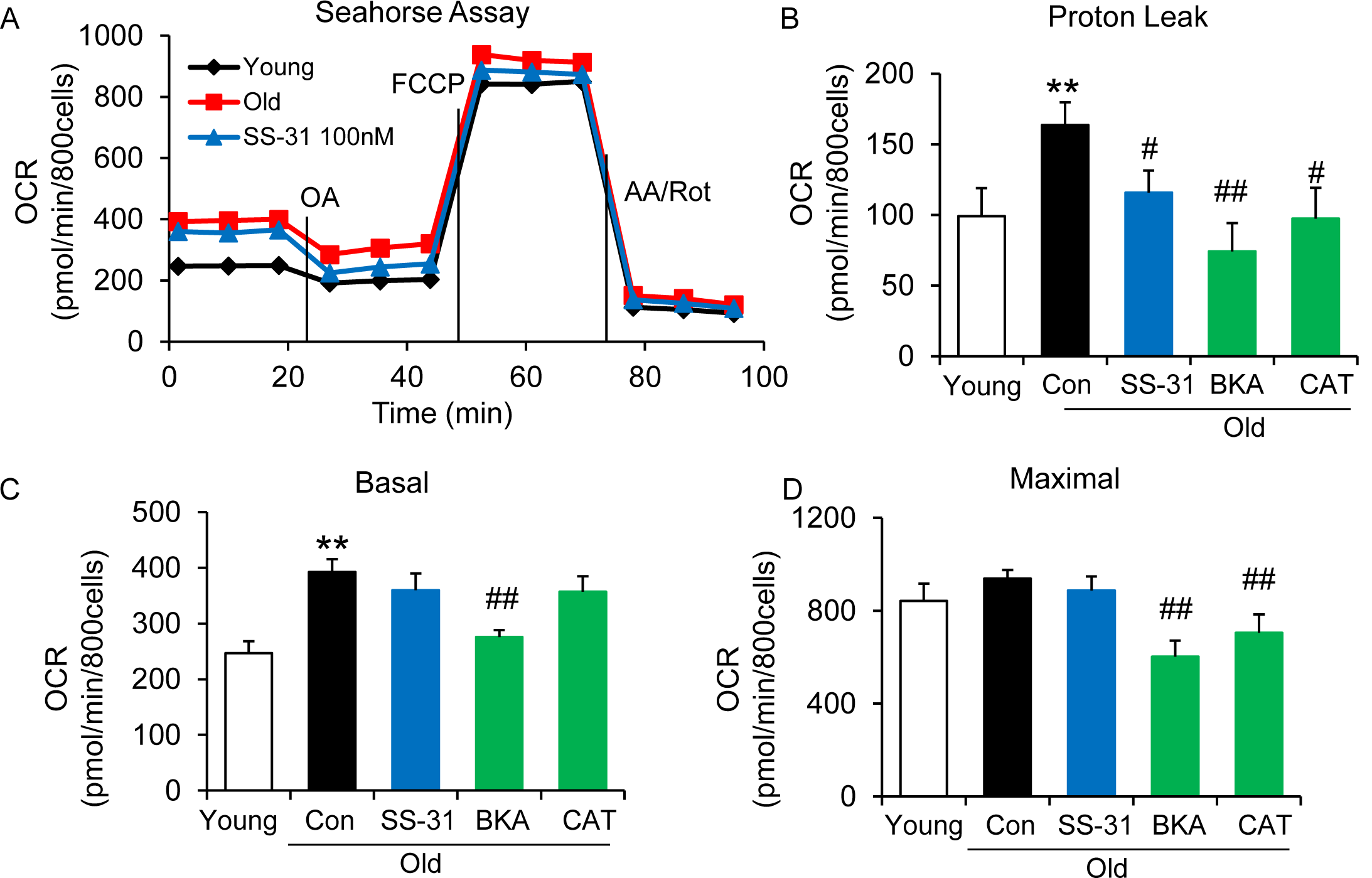
SS-31 alleviates the excessive mitochondrial proton leak of cardiomyocytes from 24 mo old mice. (A) Representative Seahorse Assay traces of cardiomyocytes isolated from untreated young and old mice, then exposed or not to 100nM SS-31 for 2 hr in vitro. Aging increased basal respiration (C), which was attributable to the augmentation of proton leak (B), but did not affect maximal respiration (D). ANT1 inhibitors BKA (10 μM) and CAT (20 μM) 2 hr treatment decreased the proton leak (B) and basal respiration (C) but also decreased maximal respiration in old cardiomyocytes. N=5-14 mice in each group; Student’s t-test was applied to determine the statistical significance. *P < 0.05, **P < 0.01 vs. young; #P < 0.05, ##P < 0.01 vs. old controls.

### SS-31 restores the resistance to external pH gradient stress in old cardiomyocytes

The evaluation of mitochondrial proton leak by Seahorse Assay is indirect, as it is based on the oxygen consumption rate. Thus, to directly investigate the reduction of mitochondrial proton leak in old cardiomyocytes by SS-31, we expressed the protein mt-cpYFP, a mitochondrial matrix-targeted pH indicator (Demaurex and Schwarzlander, 2016; Schwarzlander et al., 2012; Wang et al., 2016a; Wei-LaPierre et al., 2013) in the rat cardiomyocytes (Fig. S3). Taking advantage of the pH sensitive character of mt-cpYFP, we developed a novel protocol to evaluate mitochondrial proton leak by exposing mitochondria to a pH gradient stress in saponin permeabilized, mt-cpYFP expressing cardiomyocytes (Fig. 2A, B). The drop in mt-cpYFP 488/405 ratio is due to proton leak through the mitochondrial inner membrane into the mitochondrial matrix. To evaluate the physical properties of the mitochondrial inner membrane in the absence of mitochondrial activity, we permeabilized the cardiomyocytes in a buffer that contained no substrates, ATP, or ADP. We found that aging reduced cardiomyocyte mitochondrial resistance to a proton gradient stress (Fig. 2A, B). More importantly, we found 10 μM SS-31 treatment *in vitro* restored cadiomyocyte mitochondrial inner membrane resistance to the pH gradient stress in the aged cardiomyocytes (Fig. 2A, B). SS-31 treatment largerly prevented the decline in matrix pH of old cells after the external pH was reduced to 5.3 (Fig. 2B, C) and slowed the rate of cpYFP 488/405 change after pH 6.9 (Fig. 2D, 3C). At pH 4.5 SS-31 continued to enhance resistance to proton permeability in the treated old cells. This is unlikely to represent a biological benefit at this nonphysiologic pH, but does indicate the substantial change in physical properties of the inner membrane after interaction with SS-31 (Fig. 2B). To further evaluate the kinetics of SS-31 effect on mitochondrial proton permeability, we analyzed cpYFP fluorescence ratios at various times after exposure of the saponin treated cardiomyocytes to 10 μM SS-31. SS-31 protection of on the mitochndrial matrix proton entry became significant and near maximal after 7-10 minutes of SS-31 treatment (Fig. 2E). We examined the dose effect of SS-31 on proton permeability and found near-maximal effects at 100 nM SS-31 (Fig. 2F). In summary, this is the first direct evidence that aging increases mitochondrial inner membrane proton permeability in aged cardiomyocytes and that SS-31 protects cardiomyocytes from this proton leak.

**Fig. 2:**
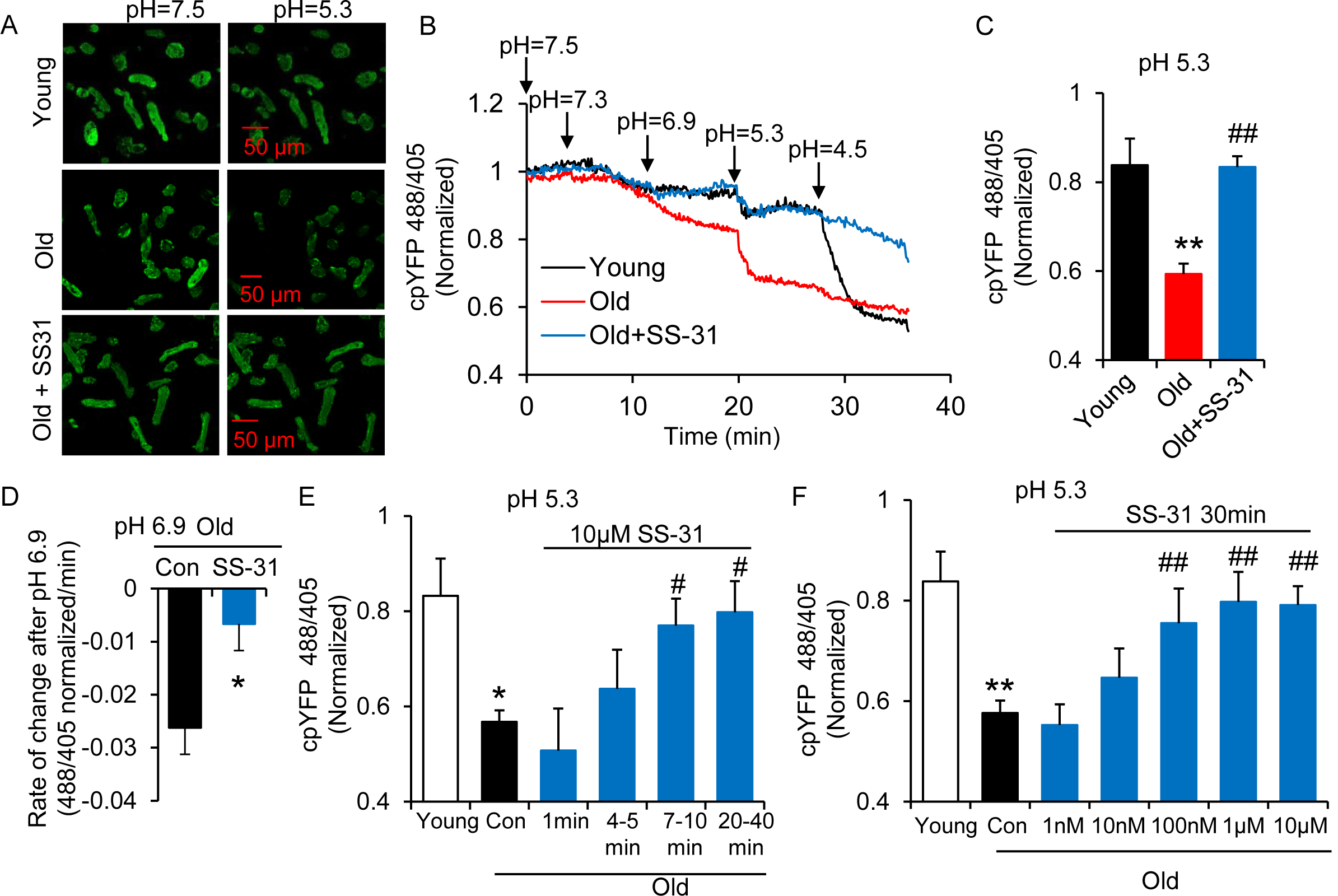
SS-31 restores the resistance of cardiomyocytes from old rat to proton entry into the mitochondrial matrix during external pH gradient stress. (A) Typical image of the effects of pH gradient stress on permeabilized rat cardiomyocyte mt-cpYFP fluorescence. Up panel, Young, middle panel, Old, lower panel, Old+SS31 (10 μM, 3 days) visualized after exposure of the cells to pH 7.5 and, later, to pH 5.3. The excitation is 488 nm and collection is at 505-730nm. (B) saponin (50 μg/ml) permeabilized cardiomyocytes expressing mt-cpYFP were exposed to progressively lower external pH. Proton permeability of old mitochondria was greater than that of young mitochondria, but preincubation of old cells with 10 μM SS-31 for 3 days enhanced the mitochondrial inner membrane resistance to the pH stress. The traces were averaged from 4-19 experiments. The arrows indicate the changes of pH. (C) Quantitation of the SS-31 treatment effect on the mitochondrial matrix cpYFP ratio at pH 5.3. The data are from 7-8 min after the pH was adjusted to 5.3. N=5-19 rats in each group. (D) SS-31 decreased the rate of cpYFP 488/405 ratio drop at pH 6.9. N=4-10 rats in each group. The rate is calculated as indicated in Figure 3C. The time dependence (N=3-4 rats in each group) (E) and dose dependence (N=3-14 rats in each group) (F) of SS-31 protection of mitochondrial resistance to pH gradient stress are shown. After cardiomyocyte permeabilization, 10 μM SS-31 was added for the times shown in or at the doses shown in (F) for 30 minutes, followed by pH stress. * P < 0.05, **P < 0.01 vs Young; # P < 0.05, ##P < 0.01 vs Old.

### ANT1 inhibitors restore resistance of old cardiomyocytes to proton leak

In search of the source of the uncoupled proton leak in the aged cells, we examined possible involvement of proton leakage through ATPase and mitochondrial uncoupling proteins (UCPs). The ATPase inhibitor Oligomycin A failed to inhibit the proton leak in pH challenged permeabilized aged cells (Fig. 3A, B). Levels of UCP2, which is the dominant isoform of UCPs in the heart, do not change with in age in hearts (Fig. S4). Genipin, an inhibitor of UCP2, showed no effect on the proton leak in permeabilized aged cells (Fig 3A, B). These results suggest that the ATPase and UCP2 may not be the source of the excess proton leak in the aged hearts.

**Fig. 3:**
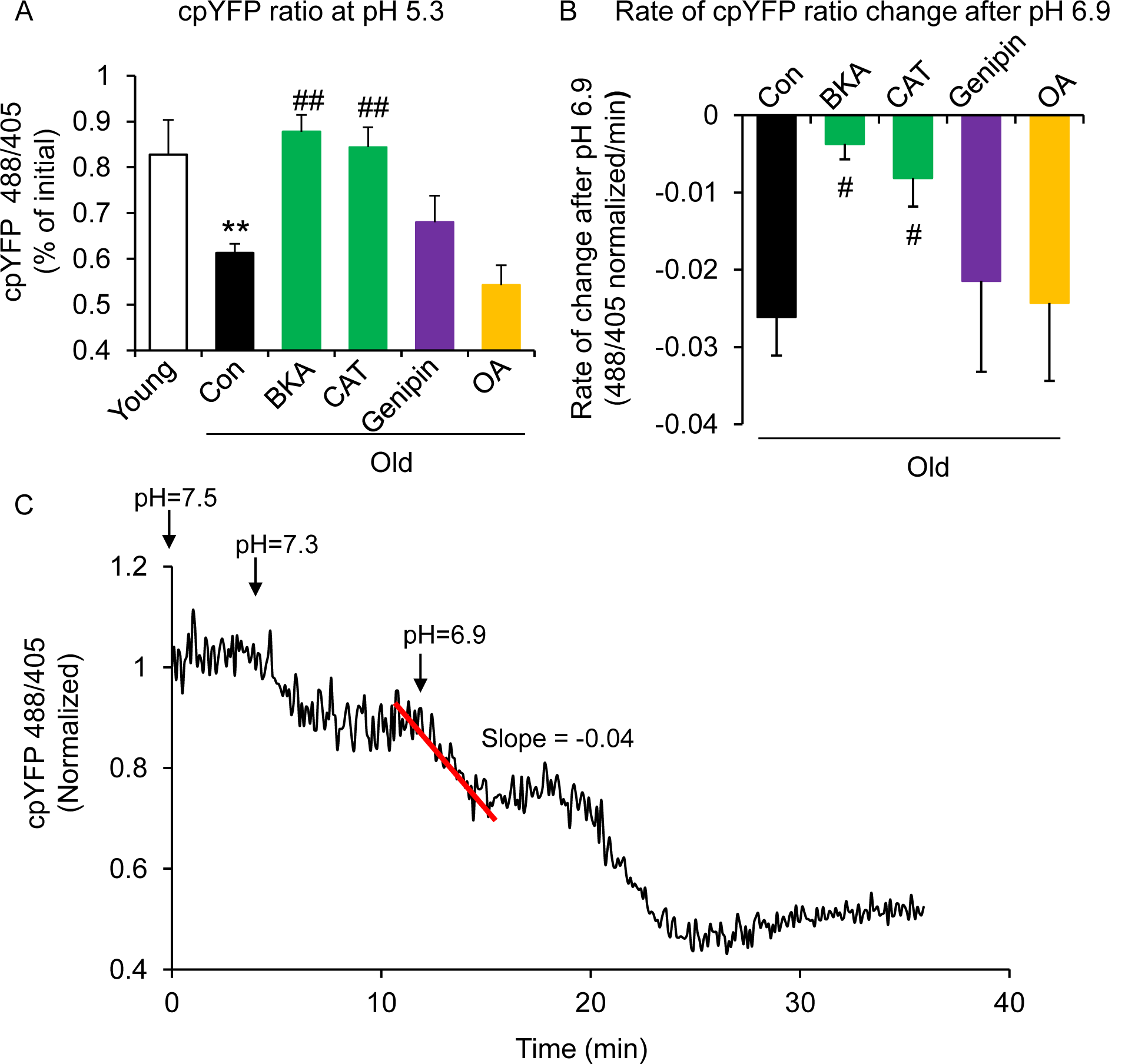
ANT1 inhibitors restore resistance to proton leak in old rat cardiomyocytes. ANT1 inhibitors 10 μM BKA and 20 μM CAT, but not 50 μM Genipin (UCP2 inhibitor) or 1 μM OA (ATPase inhibitor), protected the mitochondrial matrix from decreased pH after exposure to external pH 5.3 (N=4-19 rats in each group) (A) and reduced the rate of 488/405 decline after exposure to pH 6.9 (N=4-10 rats in each group) (B). BKA, CAT, Genipin, or OA were added immediately after the mitochondria permeabilization. *P < 0.05, **P < 0.01 vs Young, #P < 0.05, ##P < 0.01 vs Old. (C) Method of quantitation of the slope of mt-cpYFP fluorescence ratio change after permeabilized cells are exposed to external pH 6.9. The trace is from old rat cardiomyocytes.

Recently, the inner membrane protein ANT1 (also called AAC) was identified as the major site of proton leak in mitochondria of multiple tissues (Bertholet et al., 2019), and was shown to contribute to the majority of the proton leak in muscle cells (Brand et al., 2005). Treatment of old cardiomyocytes with either the ANT1 inhibitor bongkrekic acid (BKA) (Ruprecht et al., 2019) or carboxyatractyloside (CAT) (Pebay-Peyroula et al., 2003) completely suppressed the excess proton leak in the Seahorse assay, though unlike SS-31, they also decreased the maximal respiratory rate and failed to enhance the RCR (Fig 1B, C, E), which is consistent with the effect seen in the ANT triple knockout model (Karch et al., 2019). We treated permeabilized old cardiomyocytes with the ANT1 inhibitors and examined the mt-cpYFP response to an external pH gradient using the protocol described above. BKA suppressed the proton leak in old cardiomyocytes, evidenced by the preserved 488/405 ratio at pH 5.3 and a slower 488/405 ratio decrease at pH 6.9 (Fig. 3A, B). Similar inhibition was found with CAT treatment (Fig. 3A, B). Taken together, these data implicate ANT1 as the major site of proton leak in aging hearts.

### SS-31 attenuates the excessive mitochondrial flash (mitoflash) activity of aged cardiomyocytes, while normalizing membrane potential and ROS

The mitoflash (Feng et al., 2017; Hou et al., 2014; Shen et al., 2014; Wang et al., 2008; Wang et al., 2016b; Zhang et al., 2015), is triggered by nanodomain proton influx into the mitochondrial matrix (Wang et al., 2016c). Thus, we wondered whether the increased proton leak in the old cells triggered excessive mitoflash activity. We evaluated mitoflash activity in isolated young and old rat cardiomyocytes using the indicator mt-cpYFP, as established in the previous studies noted above. The mitochondrial mitoflash activity in the cells from old (26 mo) cardiomyocytes was higher than that of young (5 mo) cells (2.8 ± 0.3 in old vs 1.4 ± 0.2, /1000μm^2^/100s in young cells, n=28-88, p<0.05). Confirming this, we detected an increase in mitoflash activity in Langendorff perfused intact aged hearts from mt-cpYFP transgenic mice (Fig. S5). 1 hour treatment with SS-31 normalized the mitoflash activity in old cells to the young cell level (Fig. 4D). Moreover, the mitochondrial ANT1 inhibitors BKA and CAT showed super-suppression of the flash activity, reducing this frequency to half of that of young cells (Fig. 4D). These data support the notion that proton leak from ANT1 triggers the mitoflash in cardiomyocytes and is responsible for the excess mitoflash activity of old cells. Moreover, the mitochondrial membrane potential, which is generally lower in old cardiomyocytes (Serviddio et al., 2007), is restored to youthful levels by SS-31 treatment (Fig. 4E). Also, SS-31 reduced ROS production in the aged cardiomyocytes (Fig. 4F). Thus, the reduction of mitochondrial proton leak by SS-31 is accompanied by a more youthful membrane potential and dynamic function (mitoflash), as well as less oxidative stress.

**Fig. 4:**
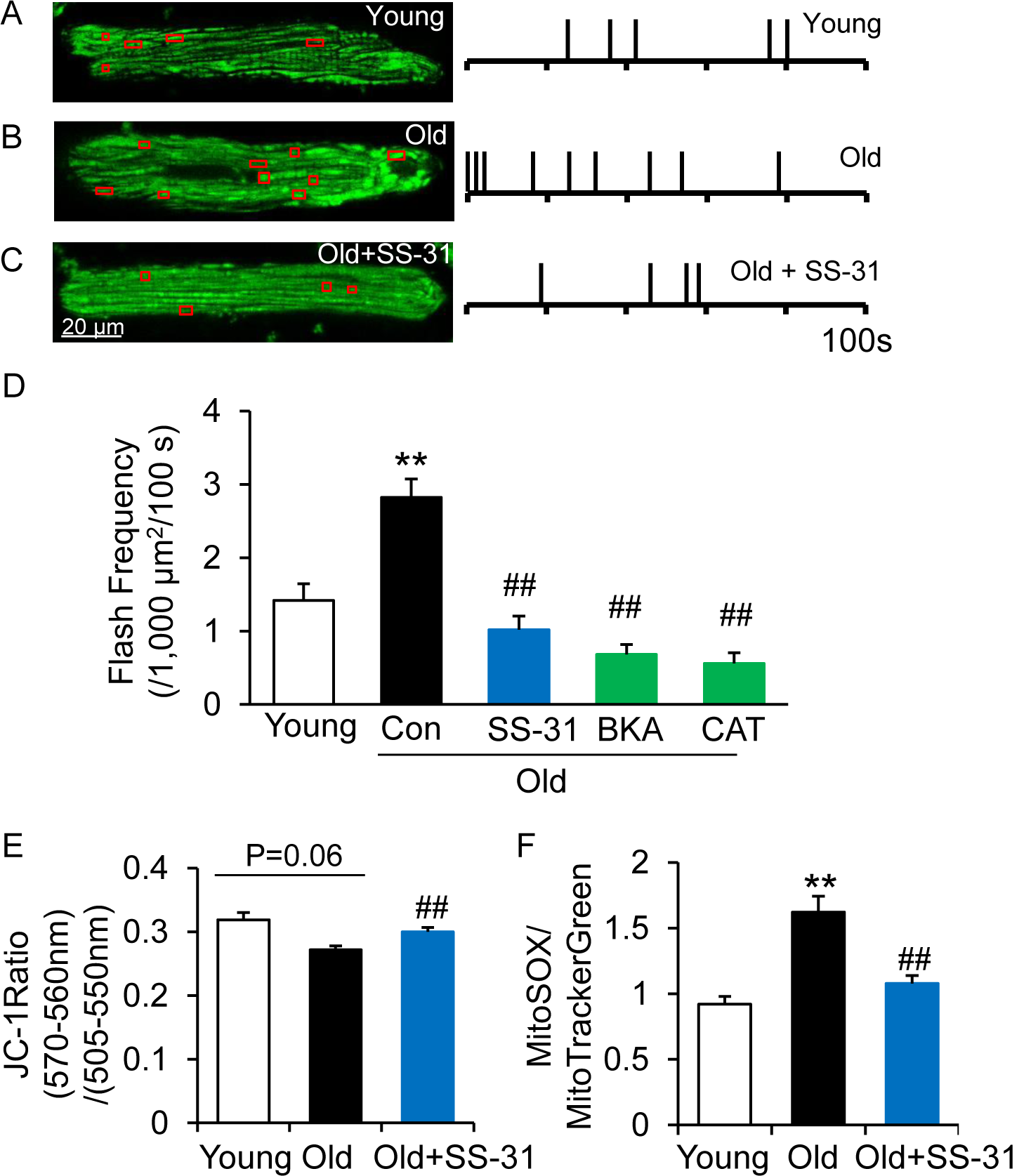
SS-31 attenuates excessive mitoflash activity in aged rat cardiomyocytes. (A-C) Mitoflash events within the regions shown in the red boxes took place at the times shown by vertical bars during the 100 sec scanning time in the representative cardiomyocytes from young (A), old (B) and old+SS-31 (C) rat hearts. (D) The rate of mitoflash activity was increased in old rat cardiomyocytes compared to young, but 1 hour SS-31, BKA (10 μM) and CAT (20 μM) treatments decreased the mitoflash frequency in old cells to or below that of young cells. N = 26-87 cells from 3-14 rats. **P < 0.05 vs young. ##P < 0.05 vs old. (E) JC-1 red to green fluorescence ratio, indicative of mitochondrial membrane potential, in cells from young and old mice and old mouse cardiomyocytes treated with SS-31 (10 μM for 12 hours). N= 84-218 cells from 3-4 mice. P = 0.06 vs young. ##P < 0.01 vs old. (F) Mitochondrial ROS production in mouse cardiomyocytes measured by the fluorescence ratio of MitoSOX (5 μM, excitation 540 nm, emission >560 nm) to Mitotracker green (200 nM, excitation 488 nm, emission 505-530 nm). N=40-84 cells from 3-5 mice. **P < 0.05 vs young. ##P < 0.05 vs old.

### SS-31 reverses increased mPTP opening in aged cardiomyocytes

Due to the close link previously established between the mitoflash and mitochondrial permeability transition pore (mPTP) opening (Hou et al., 2014), we evaluated mPTP activity by the photon-triggered mPTP opening protocol (Fig. 5A) (Zorov et al., 2000). Consistent with previous reports in isolated mitochondria (Hafner et al., 2010), we found that the time to mPTP opening is decreased in intact old cardiomyocytes (Fig. 5B). SS-31 and the ANT1 inhibitor BKA, which stabilizes the ANT1 in the m-state open towards the mitochondrial matrix, both protect the aging-increased mitochondrial mPTP opening rate (Fig. 5B), consistent with previous observations that BKA prevents the onset of the permeability transition (Halestrap et al., 1997). The ANT1 inhibitor CAT, which stabilizes ANT1 in the c-state open toward the cytosol, failed to prevent the rapid opening of the mPTP in old cells (Fig. 5B), consistent with previous observations that it facilitates mPTP opening (Halestrap et al., 1997). These data indicate that SS-31 decreases mPTP opening in old cardiomyocytes.

**Fig. 5:**
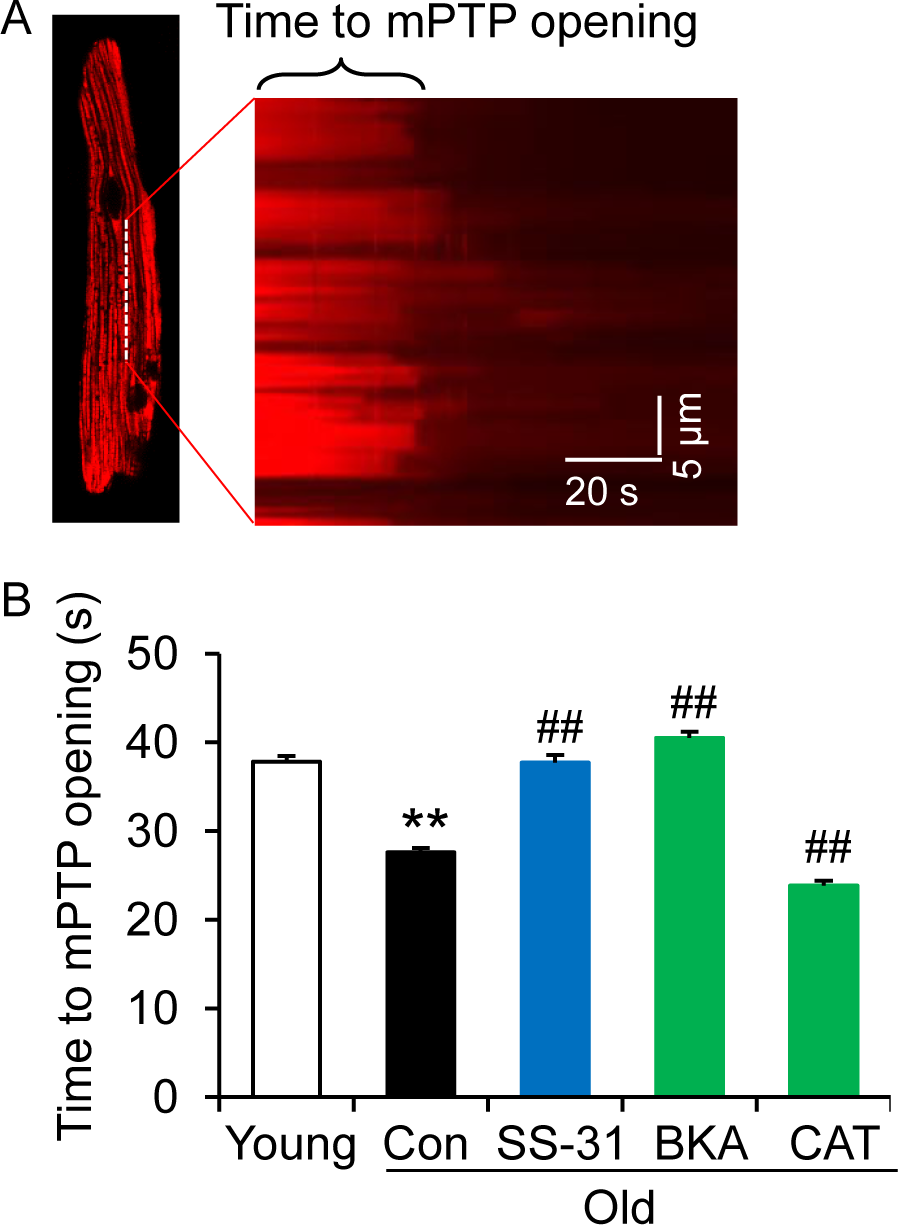
SS-31 reverses the increased speed of mPTP opening in aged mouse cardiomyocytes. (A) A typical image shows 1 Hz line-scanning photo-excitation induced mPTP opening in a cardiomyocyte loaded with the mitochondrial membrane potential (ΔΨ_m_) dye TMRM. The sudden decline of TMRM fluorescence with time (rightward) indicates mPTP opening and ΔΨ_m_ loss. (B) 1-hour SS-31 and BKA, but not CAT treatments protect the photo-excitation induced mPTP opening. Quantification of time to mPTP opening from 418-658 mitochondria from 19-32 cells isolated from 3-4 mice in each group. **P < 0.01 vs young, ##P < 0.01 vs old.

### SS-31 associates directly with ANT1 and the ATP synthasome

To further investigate the mechanism of SS-31 protection of the proton leak, we used biotinylated SS-31 to evaluate whether SS-31 directly interacts with the ANT1 protein. Hearts were disrupted by douncing, after a low speed spin to remove fragments, mitochondria were collected by high speed spin and disrupted in digitonin, to create lipid rafts containing their associated proteins, a protocol commonly used to prepare mitochondrial supercomplexes (Johnson et al., 2013). This preparation was incubated with SS-31-biotin or biotin only, followed by incubation with streptavidin beads. After washing, the bead-bound fraction was eluted with excess SS-31 and analyzed by Western blotting. Biotin-SS-31 pulled down ANT1, and free SS-31 competed with the biotin-SS-31 binding to ANT1 (Fig. 6A, B, Fig. S6). Most notably, both BKA and CAT inhibited binding of biotin-SS-31 to ANT1 (Fig. 6A, B). This competition was observed even at BKA and CAT concentrations in the tens of namomolar range (data not shown), which is consistent with their reported Kd of binding to ANT1 (Vignais et al., 1976). Biotin-SS-31 pulldown of ANT1 was not inhibited by Genipin or Oligomycin A (Fig. 6A, B). These data indicate that SS-31 associates closely with the ANT1 protein. Moreover, native gel and ATPase blot analysis showed that SS-31 stabilized the ATP synthasome, of which ANT1 and ATPase are critical members (Ko et al., 2003) (Fig. 6D, E). However, SS-31 treatment did not produce a detectable increase in mitochondrial complex proteins by Coomassie blue staining (Fig. 6C). Taken together, these data suggest that SS-31 interacts directly with ANT1 and stabilizes the ATP synthasome in old cardiomyocytes.

**Fig. 6:**
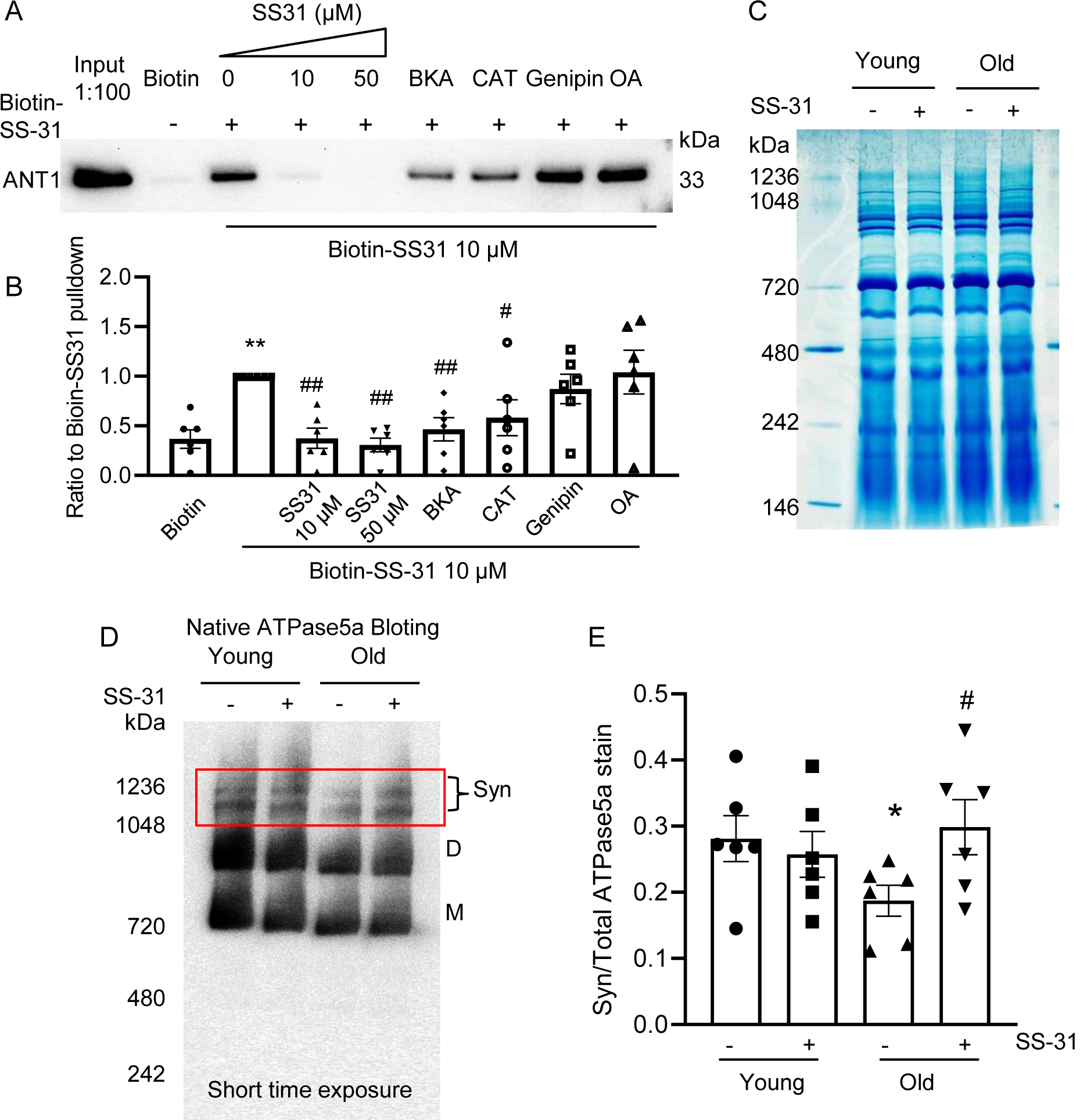
SS-31 interacts with ANT1 and stabilizes the ATP synthasome in the old mouse heart mitochondria. (A, B) Biotin-SS-31 pulldown shows the association of biotin-SS-31 to ANT1. Free SS-31 competes with this interaction, while BKA and CAT inhibit the interaction of biotin-SS-31 with ANT1. Panel A shows a representative Western blot. N=6 mice in each group. **P < 0.01 vs Biotin control, #P < 0.05, ##P < 0.01 vs Biotin-SS31 pulldown. (C) Coomassie blue staining of isolated mitochondria in a native gel. (D) Left panel is the total protein loading control for the Native gel blot. (D, E) Native Gel blotting shows that 10 μM SS-31 stabilizes the mitochondrial synthasome (Syn) in isolated mitochondria. The Syn is highlighted in the red box. The Syn and ATPase Dimer (D) and Monomer (M) were labeled using anti-ATP5A. N=6 mice in each group. *P < 0.05 vs young, #P < 0.05 vs old.

## Discussion

In this report we have shown direct evidence of increased proton leak in the aged mitochondria as a primary energetic disturbance and evidence that the increased proton entry in old cardiomyocytes takes place through ANT1. Moreover, we demonstrated that SS-31 prevents the proton entry to the mitochondrial matrix and rejuvenates mitochondrial function through direct interaction with ANT1 and stabilization of the ATP synthasome. During aging, the pathological augmented and sustained basal proton leak burdens the mitochondrial work load, resulting in a decline in respiratory efficiency. Blocking this pathological proton leak induced by aging benefits the mitochondria and the heart (Fig. 7). We suggest that the restoration of aged mitochondrial function that is conferred by SS-31 is directly attributable to this effect. However, the resulting enhancement in diastolic function is likely to require downstream changes, as the functional benefit took up to 8 weeks to reach full effect, and required post-translational modifications of contractile protein elements (Chiao et al., 2020). It is increasingly recognized that mitochondrial function, including redox status and energetics, has far-reaching effects, including epigenetic alterations and post-translational modifications (Olgar et al., 2019).

**Fig. 7:**
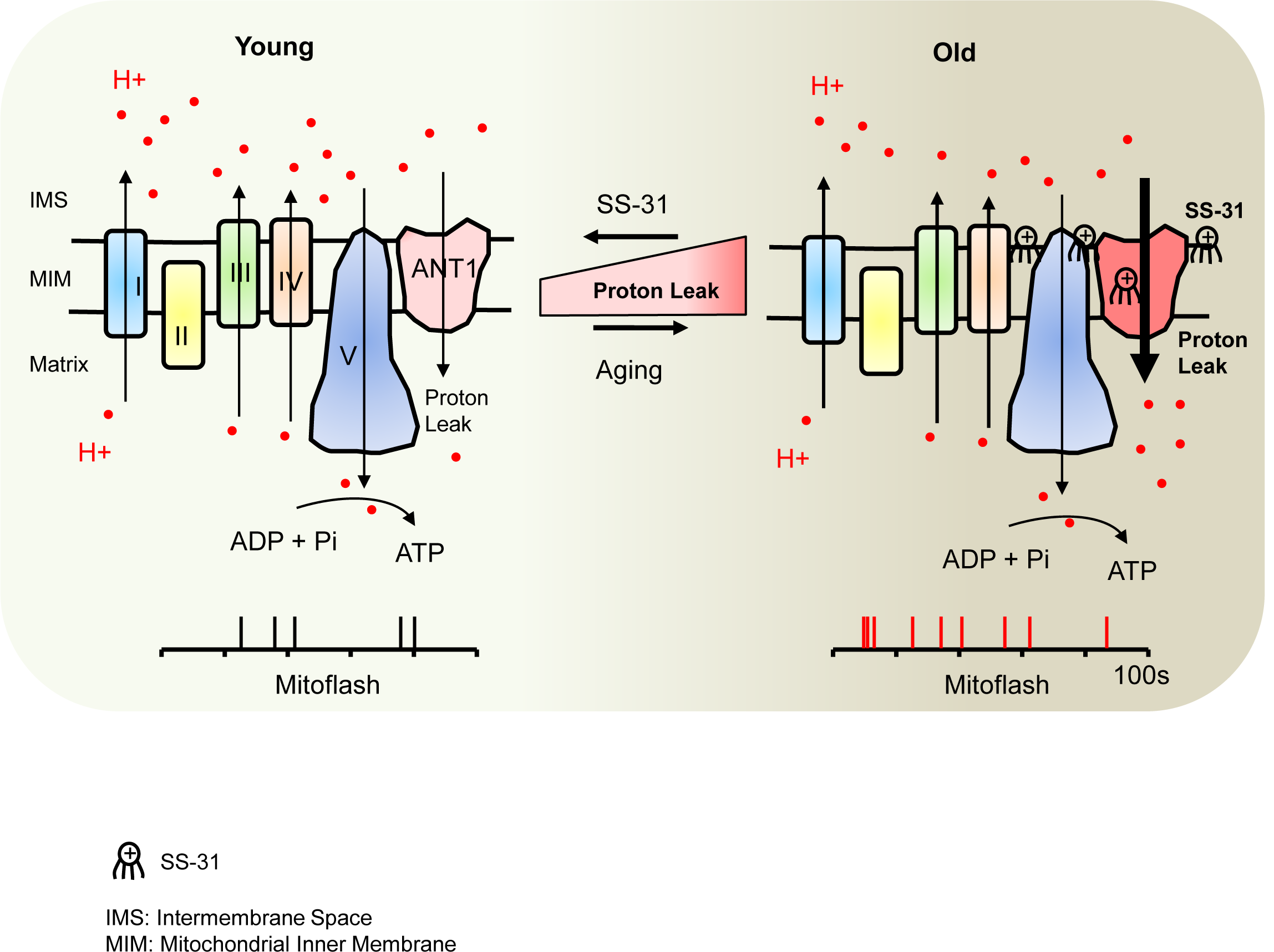
Schematic of the mechanism of SS-31 protection of proton leak and rejuvenation of mitochondrial function. Due to increased mitochondrial proton leak, the mitochondria work harder to maintain ATP production, and thus the work load is increased in the aged heart.

ANT1 appears to mediate the pathological mitochondrial proton leak in the aged mouse heart. Although an increased mitochondrial proton leak in the aged heart was previously suggested by indirect oxygen consumption measurement (Serviddio et al., 2007), the site of this augmented proton leak in aging mitochondria has remained a puzzle. We directly evaluated the proton leak using the mitochondrial matrix targeted pH indicator (mt-cpYFP) and provide evidence that implicates ANT1, the ATP/ADP translocator, as responsible for the pathologically increased proton leak in aged cardiomyocytes. This does not necessarily implicate the ADP/ATP translocase mechanism itself in the proton leak, as in the unenergetic state in which we examined the mitochondrial pH resistance, there would be no ADP/ATP transport activity. Our result is, however, supported by a recent report that proton transport is an integral function of ANT1 (Bertholet et al., 2019). Because the ANT1 protein level is not increased in the aged heart (it is, in fact, mildly but significantly decreased, Fig. S3), the aging-augmented proton leak through ANT1 must be through altered transport activity or conformational change. Both the inhibitors of BKA (locking ANT1 in m-state (Ruprecht et al., 2019)) and CAT (locking ANT1 in c-state (Pebay-Peyroula et al., 2003)) suppressed the proton leak in the aged cardiomyocytes, suggesting that constraining the conformational state in either position, or otherwise blocking the proton pore reduces ANT1 proton translocation.

Most interestingly, for the first time, we showed that a novel drug, SS-31 (elamipretide), now in clinical trials, prevents the augmented mitochondrial proton leak, rejuvenates mitochondria function and reverses aging-related cardiac dysfunction. Mechanistically, we found that SS-31 directly interacts with ANT1 and stabilizes formation of the ATP synthasome. This would seem surprising, given the prior belief that SS-31 affects mitochondria via binding to cardiolipin. However, the notion that SS-31 prevents the proton leak by direct interaction at the pore “pocket” of ANT1 is supported by recent observations based on cross-linking “interactome” mass-spectroscopy that showed that SS-31 is in intimate proximity to two lysine amino acid residues in the water filled cavity of the ANT1 protein (Chavez et al., 2020). Moreover, the cross-linking data suggested that this interaction could have structural consequences and may stabilize the m-state of ANT1 (Chavez et al., 2020). Our observation that both BKA and CAT blocked the SS-31 interaction with ANT1 suggests that SS-31 interacts with ANT1 independent of the ANT1 face to m-state (matrix facing) or c-state (cytoplasmic-facing). These results, and prior evidence of the critical role of ANT1 in mitochondrial health and function (Liu and Chen, 2013) warrant further high resolution structural study of the ANT transporter.

It has recently been shown that SS-31 alters surface electrostatic properties of the mitochondrial inner membrane (Mitchell et al., 2019). The consequences of this effect could include alteration of the channel ion gating properties of the ANT, including conformational changes secondary to enhanced supercomplex and ATP synthasome complex stability. SS-31 effects on stabilization of mitochondrial synthasome (Figure 6) could directly contribute to the enhanced efficiency of mitochondrial respiration that is seen in muscle of SS-31 treated old animals (Siegel et al., 2013) and the improvement in performance of humans with primary mitochondrial myopathy (Karaa et al., 2018).

The restoration of membrane potential by SS-31 the old mitochondria (Fig. 4E) can be attributed to the suppression of proton leak. However, SS-31 also decreased ROS production (Fig. 4F) in the aged cardiomyocytes. It is unclear if the reduced ROS production is associated with modification of the ANT1 shown here or through a parallel mechanism. Blocking this pathological proton leak induced by aging will benefit the mitochondria and the heart. This is not in conflict with the “uncoupling to survive hypothesis”, which arises from the positive correlations between increase proton leak, reduced ROS and increased lifespan (Brand, 2000). This reduced ROS production is interpreted as resulting from decreased electromotive force and consequent reduced electron leak during transport through the respiratory chain. However, SS-31 through its interaction with cardiolipin, abundant in the inner membrane, can improve the efficiency of electron transfer, especially by its known interaction with the heme group of cytochrome c (Szeto, 2014; Szeto and Birk, 2014), thereby reducing ROS production, even as the aged mitochondrial membrane potential is increased. Thus, our data support the conclusion that SS-31 interaction with multiple inner membrane proteins enhances the performance of multiple facets of respiratory mechanics.

In summary, our study reveals that ANT1 is responsible for the elevated proton leak in old cardiomyocytes and that SS-31 directly interacts with ANT1, preventing the proton leak and rejuvenating mitochondrial function in the aged cardiomyocytes. The improved mitochondrial function leads to complex secondary changes to effect enhanced diastolic function in the aged heart. These findings provide a novel insight for better understanding of the mechanisms of cardiac aging and establish the novel concept that decreasing the pathological proton leak in the aging heart restores mitochondrial function, ultimately reversing cardiac dysfunction in aging.

## Methods

### Animals

All the animal procedures were approved by the Institutional Animal Care and Use Committee at the University of Washington and conform to the NIH guidelines (Guide for the care and use of laboratory animals). Young (4-6 month-old) and aged (24-26 month-old) C57BL/6 mice (Charles River colony) and F344 rats (25-30 month-old) were obtained from the National Institute of Aging Rodent Resource. The mt-cpYFP transgenic C57BL/6 mice were housed until reaching the age described.

### Isolation of adult mouse and rat cardiomyocytes

Single ventricular myocytes were enzymatically isolated from mouse and rat hearts as described previously (Zhang et al., 2013; Zhang et al., 2017). The rod shaped cardiomyocytes were collected by allowing cells settle down and adhere to laminin coated 24 well Seahorse plates for intact cell oxygen consumption test or to glass coverslips for confocal imaging.

### Seahorse Assay

The XF24e Extracellular Flux Analyzer (Seahorse Bioscience) was used for measuring oxygen consumption in intact cardiomyocyte, with XF assay medium containing 5 mM glucose and 1 mM pyruvate. Oligomycin A (OA, 2.5 μM), carbonyl cyanide-p-trifluoromethoxyphenylhydrazone (FCCP,1 μM), and antimycin A (AA, 2.5 μM) plus 1 μM rotenone (Rot) were added in three sequential injections. The RCR was measured or calculated by maximal respiration divided by basal respiration.

### Confocal imaging

We used a Zeiss 510 (Zeiss, Germany) or Leica SP8 (Leica, Germany) for confocal imaging at room temperature. The cells were placed in modified Tyrode’s solution (in mM: 138 NaCl, 0.5 KCl, 20 HEPES, 1.2 MgSO_4_, 1.2 KH_2_PO_4_, 5 Glucose, 1 CaCl_2_, pH 7.4). For mitochondrial flashes, mt-cpYFP expressing cells were exposed to alternating excitation at 405 and 488 nm and emission collected at >505 nm. Time-lapse 2D images were collected at a rate of 1 s per frame. For mitochondrial superoxide quantitation, we used the ratio of MitoSOX Red (5 μM, excited at 540 nm with emission collected at > 560 nm) to mitoTracker Green (200 nM, excited at 488 nm and emission collected at 505-530 nm). For mitochondrial membrane potential measurement, JC-1 was excited at 488 nm and emission collected at 510-545 nm and 570-650 nm. For photon triggered mPTP opening, the cells were loaded with 120 nM Tetramethylrhodamine methyl ester (TMRM) and line scanned at 1 Hz as described previously (Zorov et al., 2000).

### Cell permeabilization and pH stress

Rat cardiomyocytes were cultured with mt-cpYFP adenovirus (Wang et al., 2016b) for 3 days in M199 medium. After incubation in Ca^2+^-free Tyrode’s solution for 30 min, the medium was changed to a solution of 100 mM potassium aspartate, 20 mM KCl, 10 mM glutathione, 10 mM KH_2_PO_4_, 0.1 mM EGTA, 8% dextran 40,000, pH 7.5, with 50 μg/ml saponin for 30 s and then maintained in saponin-free internal solution (Lukyanenko and Gyorke, 1999). The pH of the solution containing the permeabilized cells was then progressively lowered by addition of HCl in quantities previously titrated to result in pH 7.3, 6.9, 5.3, and 4.5, with 8 min between each step. The permeabilized cells were excited using same settings as for mt-cpYFP above, but using a time-lapse of 6 s per frame. The ratio of emission fluorescence at 488 nm from 405 nm excitation indicated the mitochondrial pH change (Wei-LaPierre et al., 2013) and was normalized to a starting (pH 7.5) arbitrary value of 1.0, so as to normalize differences due to variability of the intensity of laser excitation and emission collection between different experiments.

### Western blots

Heart tissue was lysed with RIPA buffer containing a protease inhibitor cocktail (Chiao et al., 2016). Protein samples were denatured and separated via NuPAGE Bis-Tris gel, and transferred to PVDF membranes. The blots were probed with primary antibodies: ANT1 (Abcam, ab102032, 1:3000), UCP2 (Cell signaling technology, 89326S, 1:2000) followed by appropriate secondary antibodies.

### Biotin-SS-31 pulldown and Blot Analysis

Hearts were chunked and dounce homogenized in mitochondrial isolation buffer (MIB, in mM: 300 sucrose, 10 Na-HEPES, 0.200 EDTA, pH 7.4) and centrifuged at 800 g for 10 min. The supernatants were centrifuged at 8000 g for 15 min to purify mitochondria. Digitonin was added to the mitochondria at a ratio of Digitonin : protein = 6 :1 to break down the membrane system. Treatment drugs were added 30 min before addition of 10 μM biotin-SS-31 (Biotin-D-Arg-dimethyl Tyr-Lys-Phe-NH2) or biotin control (Thermo, B20656). Streptavidin Agarose beads (Thermo, 20349) were added and incubated 2 hr at room temperature. The beads were washed with MIB 3 times and then eluted by 50 μM SS-31. The eluates were boiled with LDS protein loading buffer (Thermo, NP0008) and loaded on NuPAGE for gel electrophoresis and Western blotting with antibody to ANT1 (Abcam, ab102032, 1:3000). In some experiments, after electrophoresis, gels were silver stained using a Pierce Silver Stain Kit (Thermo, #24612).

### Native coomassie blue staining and blotting

Mitochondria from mouse hearts were isolated as described previously (Marcu et al., 2012). Mitochondria (100 µg) were solubilized in 4x NativePAGE Sample Buffer containing 5% digitonin and 5% coomassie blue G-250. The samples were loaded on NativePAGE Novex 3-12% Gel and run at 100 V for 1 hr, then at 300 V for 2 hr. For coomassie blue staining, gels were stained with 0.1 % Coomassie Brilliant Blue overnight and destained with destaining solution (H_2_O: Methanol: Acetic Acid = 5:4:1) 5 times at 20 min intervals. For native blotting, gels were transferred to PVDF membranes at 25 V in 4 °C overnight and incubated with ATP5a antibody (Abcam, ab14748, 1:3000), followed by anti-mouse secondary antibody.

### Perfused mouse heart confocal imaging

mt-cpYFP transgenic mice were anesthetized with pentobarbital (150mg/kg). The heart was removed, cannulated via the ascending aorta, and put on a modified perfusion system and in a custom made chamber on the confocal stage as previously reported (Zhang et al., 2018; Zhang et al., 2017). The perfusion was maintained under a constant flow (∼2 mL/min) with O_2_/CO_2_-bubbled KHB solution (in mM: 118 NaCl, 0.5 EDTA, 10 D-glucose, 5.3 KCl, 1.2 MgCl_2_, 25 NaHCO_3_, 0.5 Pyruvate, and 2 CaCl_2_, pH 7.4) at 37 °C. To minimize motion artifact during imaging, 10 uM (-)-Blebbistatin (Toronto Research Chemicals) was included. During imaging, the left ventricle was gently pressed to further suppress motion artifact. Mitoflashes were imaged using the procedure described above.

## Data statistics

Data are shown as mean ± SEM. Student’s t-test was applied to determine the statistical significance. P < 0.05 was considered statistically significant.

## Acknowledgements

We thank Drs. Mariya Sweetwyne, Ying Ann Chiao, Martin Brand, Michael MacCoss and Gaomin Feng for technical support and helpful discussions and the services of the W. M. Keck Microscopy Center at the University of Washington. SS-31 (elamipretide) was kindly provided by Stealth Biotherapeutics (Newton MA).

This work was supported by NIA P01AG001751 and R56AG055114 to P.S.R., HL114760, HL137266 and AHA 18EIA33900041 to W. W, and a Glenn Foundation for Medical Research Postdoctoral Fellowship and AHA 19CDA34660311 to H.Z.

## Author contributions

Conceptualization and experimental design: H Zhang, N.N Alder and P.S. Rabinovitch; investigation, analysis, and visualization: H Zhang; writing of the original draft: H Zhang, P.S. Rabinovitch; review and editing of the draft: N.N Alder, W Wang, H Szeto, D. J. Marcinek.

---

**Supplemental Fig. 1:**
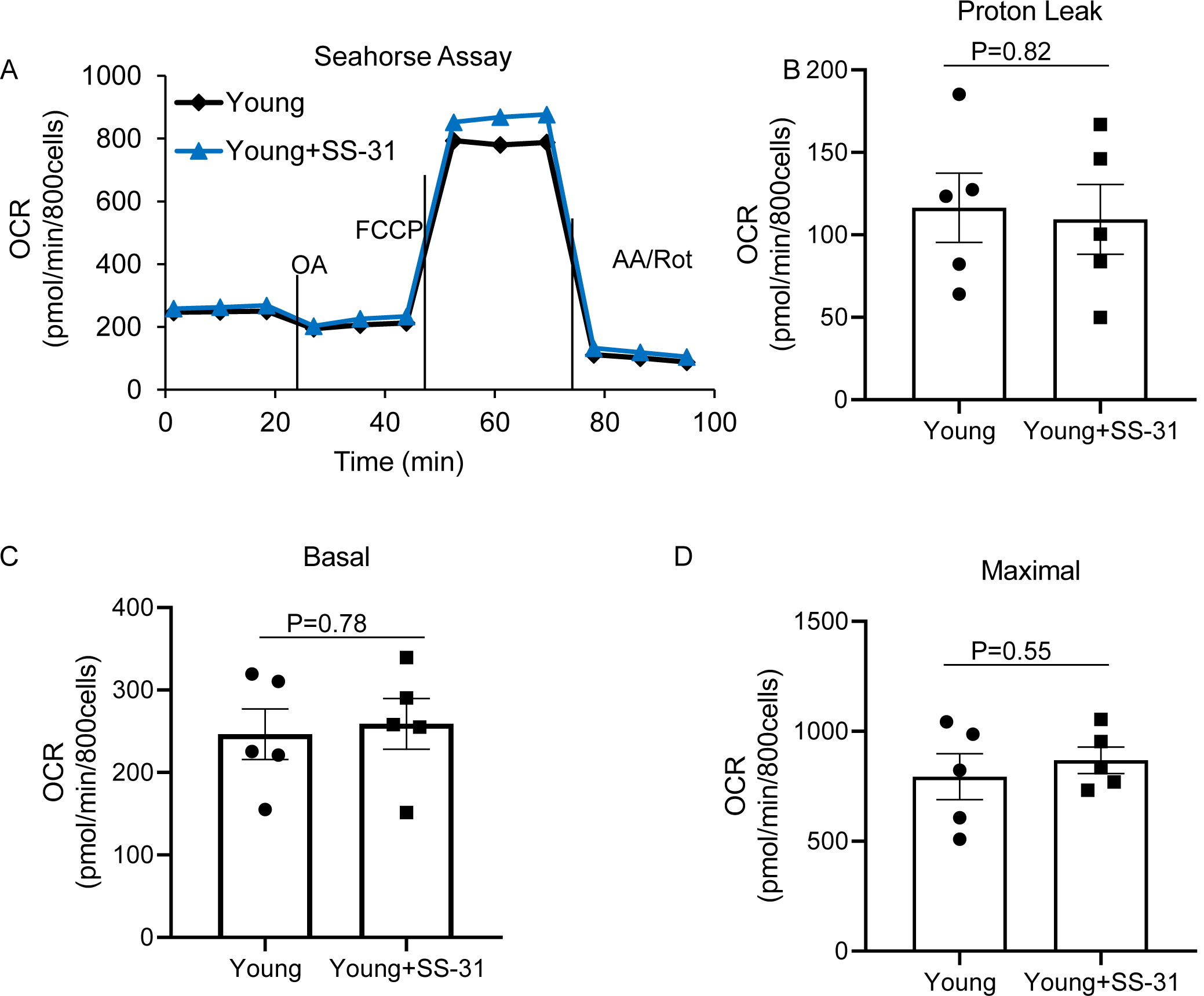
SS-31 has a minor effect on mitochondrial respiration in the young cardiomyocytes. None of the differences are significant. N=5 mice in each group.

**Supplemental Fig. 2:**
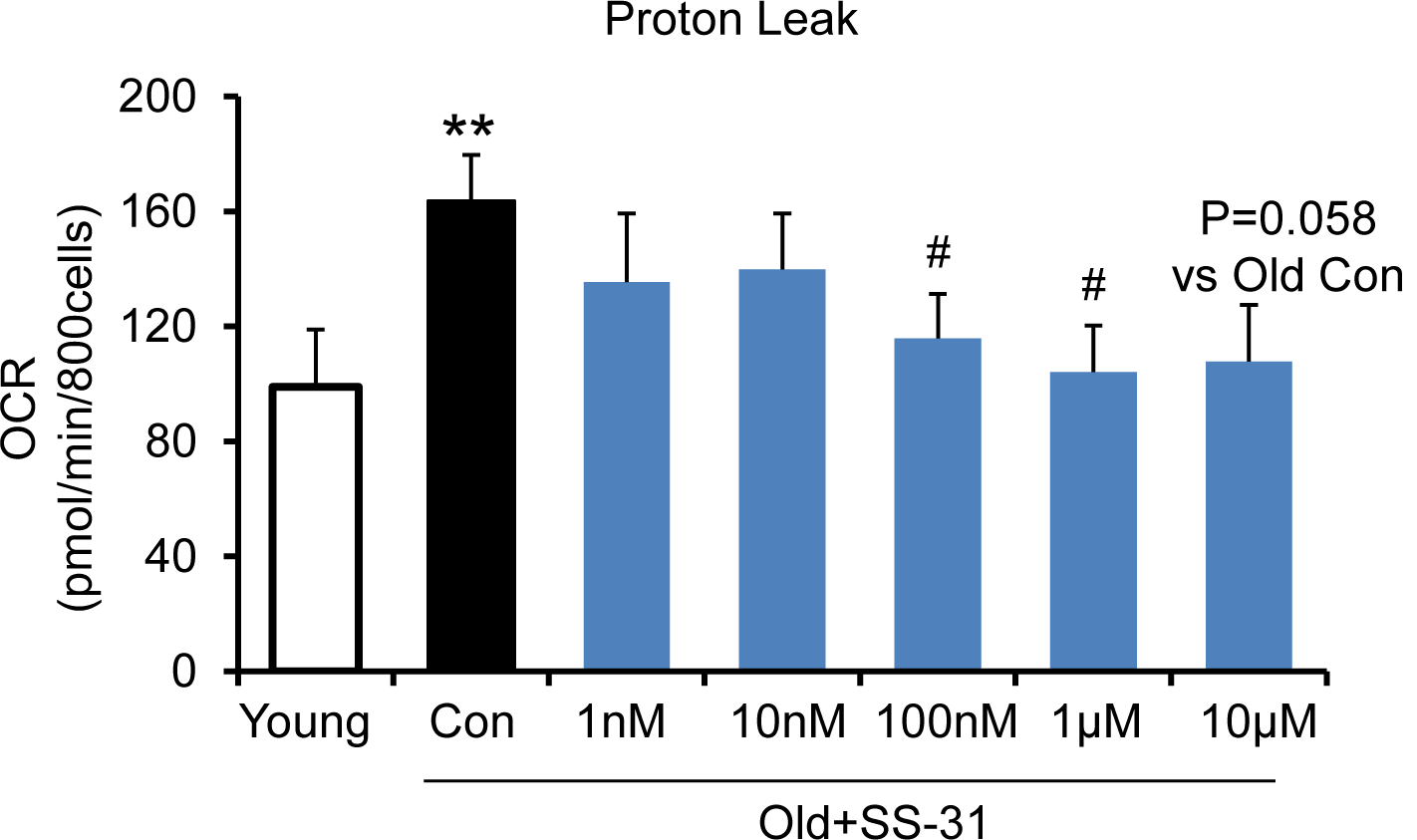
SS-31 reaches its inhibitory effect on proton leak suppression at low concentrations. SS-31 decreased the mitochondrial proton leak. N=6-14 mice in each group **P < 0.01 vs. young, #P < 0.05 vs. old controls.

**Supplemental Fig. 3:**
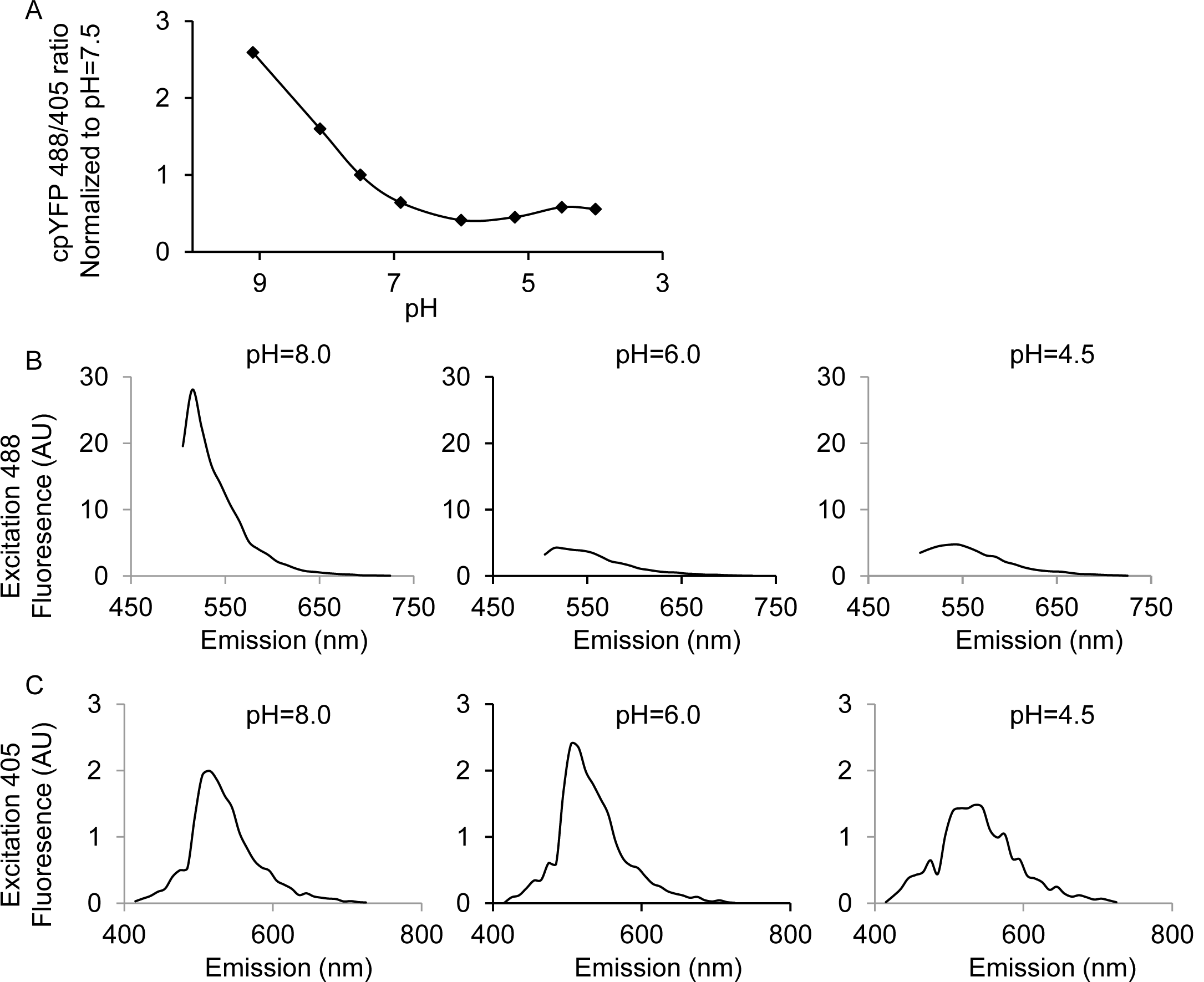
pH calibration of mt-cpYFP in adult cardiac myocytes with Nigericin. 10 μM Nigericin was added to the mitochondrial permeabilization buffer. (A) shows the ratio of 488/405 (normalized to that of the value at pH=7.5) as the pH is gradually lowered. The ratio of 488/405 reaches the lowest point at pH 6.0, followed by a small increase starting at pH 5.5, consistent with the known pH response of this dye (Wei-LaPierre et al., 2013). (B) shows the emission spectra of mt-cpYFP pH 8.0, 6.0 and 4.5 when excited at 488 nM (C) shows the emission spectra of mt-cpYFP pH 8.0, 6.0 and 4.5 when excited at 405nm. The 488 nm excitation is sensitive to the pH change but the 405 nm excitation is much less pH dependent.

**Supplemental Fig. 4:**
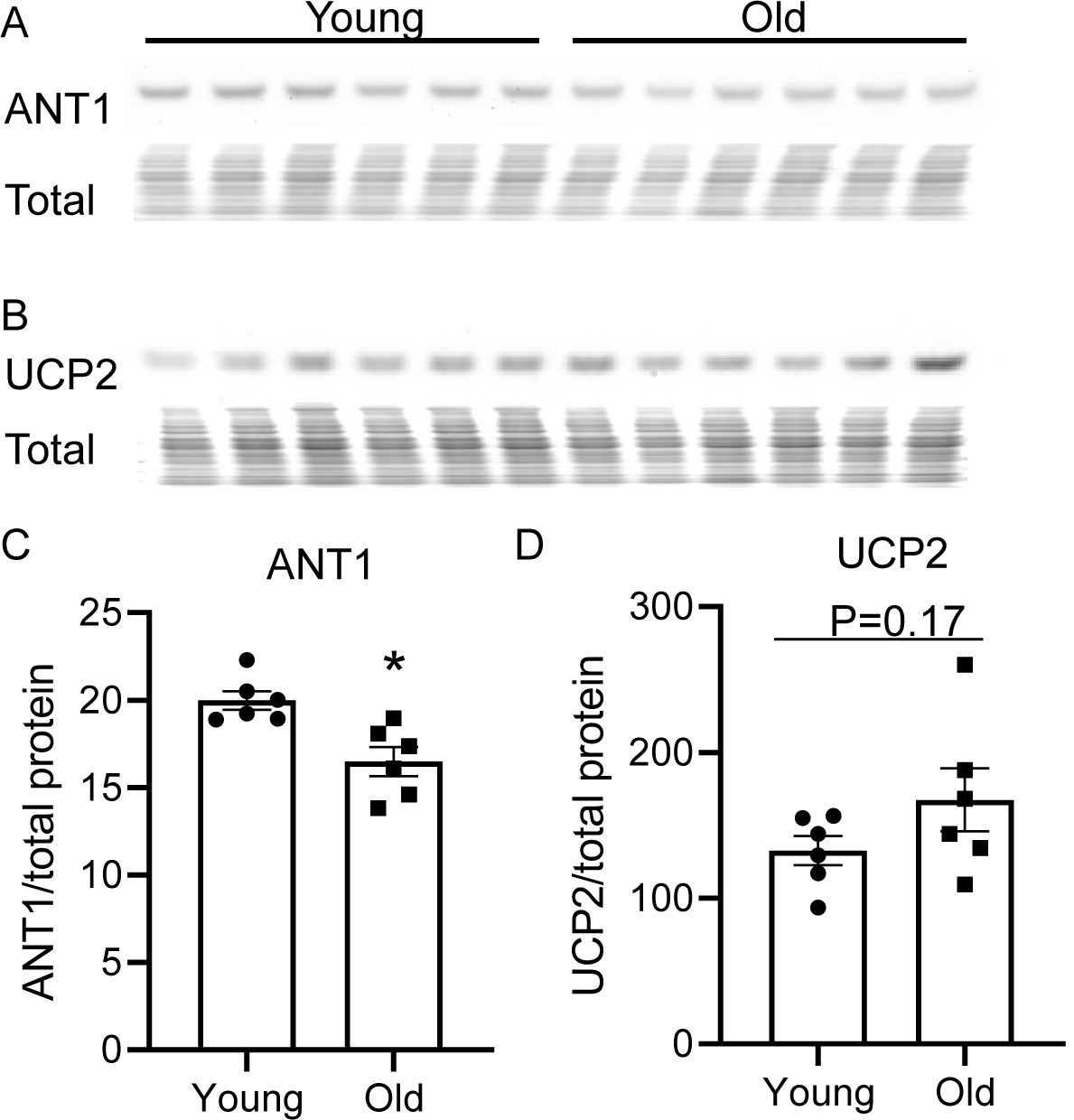
Aging effect on ANT1 and UCP2 cardiac protein abundance. Aging slightly, but significantly decreases ANT1 levels in the mice heart. Western blot of ANT1 (A, C) and UCP2 (B, D). N=6 mice in each group. *P < 0.05 vs young. The total protein loading control was stained with MemCode Reversible Protein Stain Kit for Polyvinylidene difluoride Membranes (Pierce, Rockford, IL).

**Supplemental Fig. 5:**
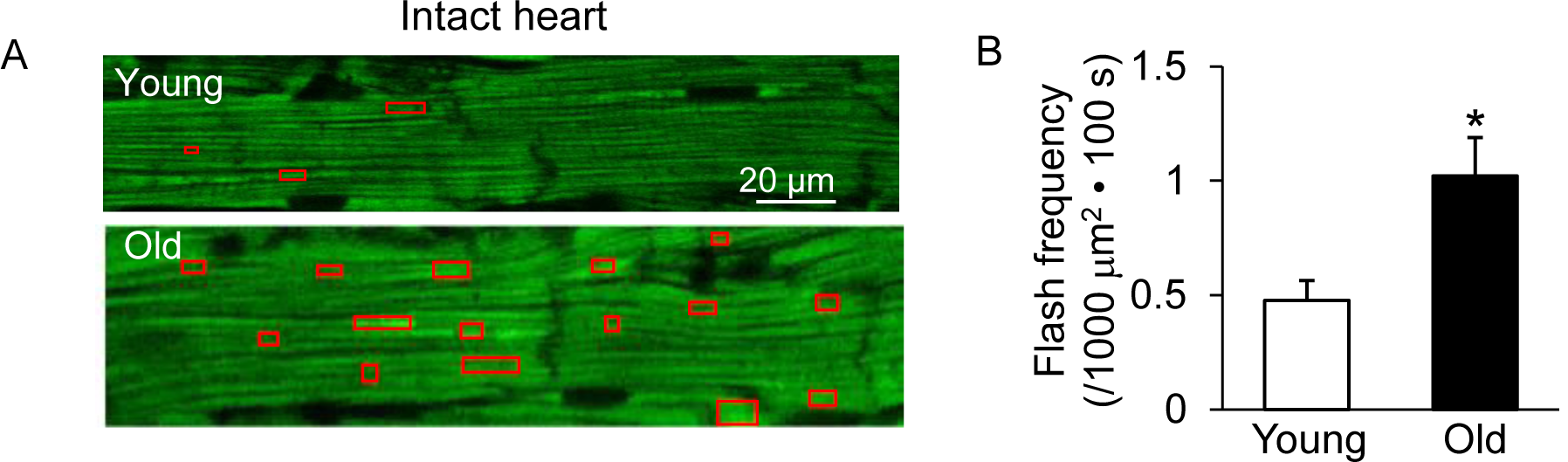
Increased mitochondrial flash activity in the intact perfused mouse aged heart. (A) Typical images from the young (upper panel) and old (lower panel) mice hearts. (B) Statistical analysis of mitoflash frequency in the analyzed regions indicated by the red boxes in panel A. Young: N=18 regions from 3 mice. Old: N=16 regions from 4 mice. *P < 0.05 vs Young.

**Supplemental Fig. 6:**
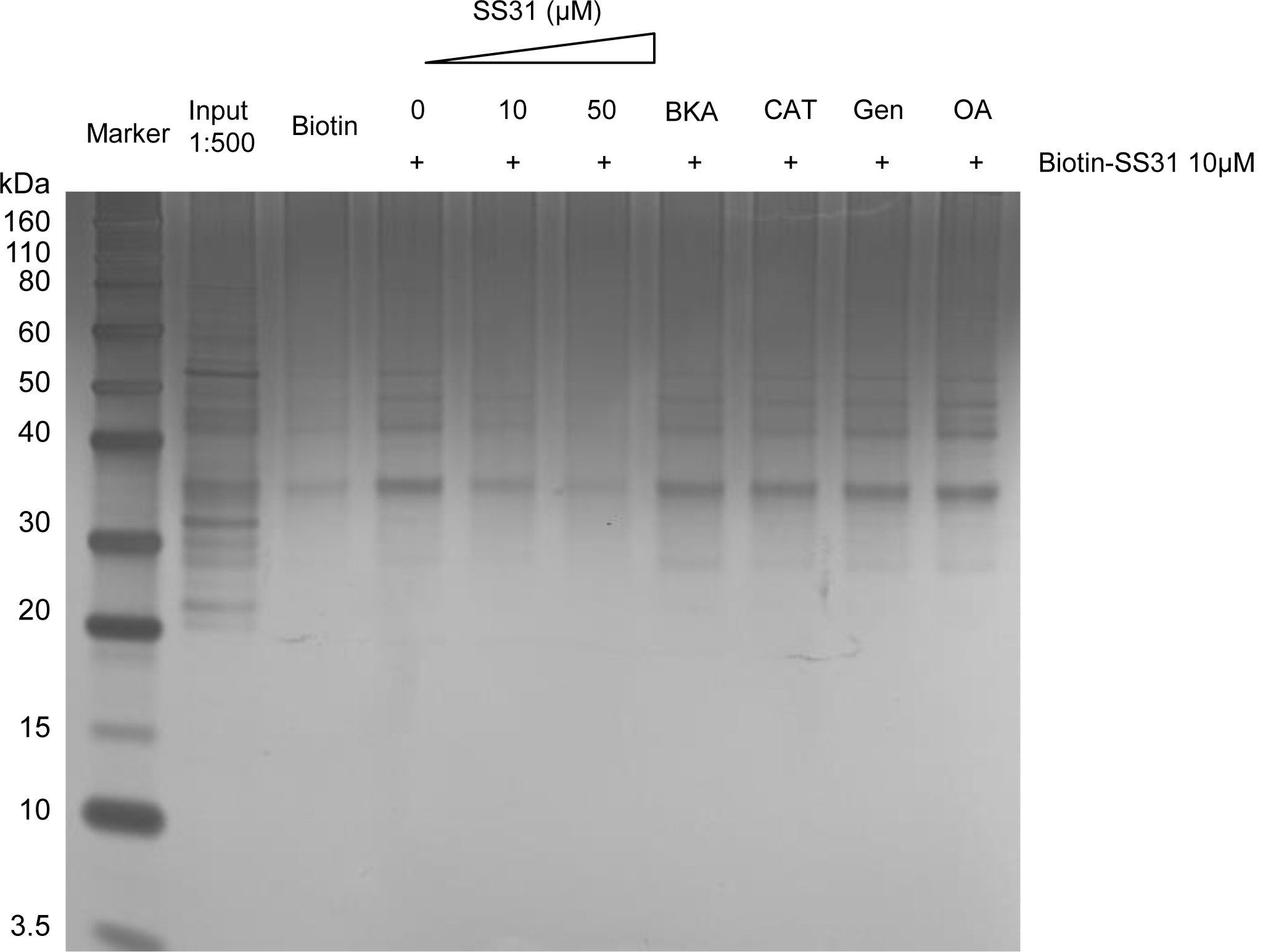
Silver staining of the Biotin-SS-31 pulldown of Figure 6A from the old mouse heart mitochondria. The great majority of bands present in the input are not present in the biotin-SS-31 pulldown; those that are show suppression by SS-31 competition.

